# Aspirin Synergizes with Regorafenib to Reduce Growth of Colorectal Cancer

**DOI:** 10.1101/2022.02.11.480021

**Authors:** Chang Su, Lochlan J Fennell, Catherine E Bond, Alexandra M Kane, Genevieve Kerr, Thierry Jardé, Diana Micati, Rebekah M Engel, Wing Hei Chan, Sara Hlavca, Stuart Archer, Paul J McMurrick, Heinz Hammerlindl, Fayth Lim, Basit Salik, Diane M McKeone, Gunter Hartel, Jennifer Borowsky, Rahul Ladwa, Barbara A Leggett, Helmut Schaider, Helen E Abud, Glen M Boyle, Matthew E Burge, Vicki LJ Whitehall

**Author notes:** Corresponding author: A/Prof Vicki Whitehall, QIMR Berghofer Medical Research Institute, 300 Herston Road, Herston, Queensland, 4006, Australia.

## Abstract

**Purpose:** Regorafenib is a multi-kinase inhibitor approved for refractory metastatic colorectal cancer. Previous studies have suggested that combining kinase inhibitors with aspirin may improve patient outcomes. We aimed to determine the effects of aspirin and regorafenib combination treatment in preclinical models of colorectal cancer.

**Experimental Design:** SW480, RKO and LIM1215 colorectal cancer cell lines were treated with aspirin and regorafenib to determine effects on proliferation and cytotoxicity. RNA sequencing and Western blotting were performed to explore underlying molecular effects. Aspirin and regorafenib combination treatment was also tested using organoids derived from three human colorectal cancer tissue specimens. For the *in vivo* study, SW480-derived tumors were established in athymic mice. Tumor volume was measured during treatment with aspirin and regorafenib, followed by immunohistochemical staining for markers of proliferation and apoptosis.

**Results:** Aspirin and regorafenib synergistically inhibited proliferation of colorectal cancer cell lines and patient-derived organoids, irrespective of *KRAS* or *BRAF* mutation status. This was associated with inhibition of the PI3K-Akt-mTOR pathway and activation of the AMPK pathway. Aspirin and regorafenib effectively inhibited growth of microsatellite stable *KRAS*-mutant SW480-derived tumors *in vivo*. Immunohistochemical staining for Ki67 and cleaved caspase 3 showed that combination treatment elicited a synergistic anti-proliferative effect, in addition to a pro-apoptotic effect that was driven by regorafenib.

**Conclusions:** Aspirin and regorafenib demonstrate synergistic anti-proliferative effects in preclinical models of colorectal cancer. This suggests that combining regorafenib with aspirin may be an improved treatment strategy for patients with refractory metastatic colorectal cancer.

## Introduction

Colorectal cancer (CRC) is the second leading cause of cancer-related mortality worldwide (1). Approximately 25% of patients present with metastatic disease at initial diagnosis, with another 25% estimated to progress to metastatic disease after treatment of early stage disease (2). Despite the availability of several chemotherapy and targeted therapy options, the median overall survival of patients with metastatic CRC is only 30 months (2), representing an unmet need for improved treatment strategies.

Regorafenib is a multi-kinase inhibitor approved as a last-line option for patients with metastatic CRC who no longer respond to combination chemotherapies, angiogenesis inhibitors or epidermal growth factor receptor (EGFR) inhibitors (3). Its use is indicated in refractory metastatic CRC irrespective of common molecular markers such as *KRAS* or *BRAF* mutation, or microsatellite instability (MSI) (3). Regorafenib inhibits B-Raf and c-Raf in the mitogen-activated protein kinase (MAPK) pathway, including B-Raf V600E activating mutation (4, 5). Regorafenib also targets various receptor tyrosine kinases including c-Kit, Ret, angiopoietin receptor Tie2, fibroblast growth factor receptors (FGFR) and vascular endothelial growth factor receptors (VEGFR) (4, 5). Several *in vitro* and *in vivo* studies of CRC have shown that regorafenib elicits anti-proliferative, pro-apoptotic and anti-angiogenic effects (4–9). However, given the broad spectrum of kinase inhibition, the molecular mechanisms underlying these effects are complex and not well understood.

Clinical studies of refractory CRC have shown that regorafenib monotherapy offers only a modest, albeit statistically significant, improvement in median overall survival of 1 to 2 months compared to best supportive care alone (10, 11). Additionally, around 70% of patients experience treatment-emergent adverse effects that necessitate dose reduction or interruption (12). Combining regorafenib with other forms of therapy may enhance anti-tumor activity, and if this can be achieved with lower regorafenib doses, may also reduce treatment-related toxicity (13).

The use of aspirin after CRC diagnosis is associated with reduced CRC-specific mortality and reduced risk of metastases (14, 15), particularly in the presence of activating *PIK3CA* mutations (16). In the refractory metastatic setting, Giampieri *et al.* found that incidental use of aspirin for cardiovascular indications is associated with improved overall and progression-free survival in CRC patients receiving last-line treatment with capecitabine followed by regorafenib (17). Additionally, Casadei-Gardini *et al.* found an association between aspirin use and improved overall survival in patients receiving last-line treatment with regorafenib for refractory hepatocellular carcinoma (18). While these observational studies suggest a role for aspirin in improving outcomes for patients receiving regorafenib in the refractory metastatic setting, combination treatment with aspirin and regorafenib has not yet been tested in pre-clinical studies of CRC or any other cancer type.

Recent pre-clinical studies have found that aspirin and the regorafenib analogue, sorafenib, interact synergistically to inhibit proliferation of CRC cells *in vitro*, irrespective of *KRAS* or *BRAF* mutation status (19, 20). We and others have demonstrated that combination treatment with aspirin and sorafenib inhibits proliferation and induces apoptosis of melanoma, lung carcinoma and hepatocellular carcinoma cells *in vitro*, and reduces growth of cell line-derived xenografts *in vivo* (21–23). The mechanisms underlying these effects are unclear but may involve activation of the AMP-activated kinase (AMPK) pathway, which mediates inhibition of mammalian target of rapamycin (mTOR) (21, 22). Modulation of the MAPK pathway may also play a role. However, both inhibition and activation of the MAPK pathway have been reported following aspirin and sorafenib combination treatment (20, 21). Additionally, sorafenib and regorafenib have distinct kinase inhibition profiles (24). The extent to which findings from studies of aspirin and sorafenib combination treatment apply to aspirin and regorafenib combination treatment is unclear, hence the need to study the combination of aspirin and regorafenib specifically. Here we show that aspirin and regorafenib synergistically inhibit proliferation of CRC cells *in vitro* and *in vivo*, providing a rationale for investigating aspirin and regorafenib combination therapy in patients with refractory metastatic CRC.

## Materials and Methods

### Cell Culture and Treatment

SW480 (American Type Culture Collection Cat# CCL-228, RRID: CVCL_0546), RKO (American Type Culture Collection Cat# CRL-2577, RRID: CVCL_HE15) and LIM1215 (CellBank Australia Cat# CBA-0161, RRID: CVCL_A4IG) cell lines, representing different molecular phenotypes of CRC (Supplementary Table S1) (25–28), were cultured in phenol red-free RPMI-1640 medium (Sigma-Aldrich Cat# R8755) supplemented with 10% fetal bovine serum (ThermoFisher Cat# 10099), 100 units/mL penicillin and 100 μg/mL streptomycin (ThermoFisher Cat# 15140). SW480 is *KRAS*-mutant (G12V), RKO is *BRAF*-mutant (V600E), and LIM1215 is *RAS/RAF*-wildtype (25, 26). Cells were maintained in a humidified 37°C incubator containing 5% CO_2_. All cell lines were routinely authenticated using short tandem repeat profiling and verified to be free of mycoplasma using the MycoAlert mycoplasma detection kit (Lonza Cat# LT07-418), by the QIMR Berghofer Medical Research Institute analytical facility (Brisbane, Australia).

### Cell Viability

Cells were harvested for experiments using 0.25% trypsin-EDTA (ThermoFisher Cat# 25200) and viable cells were counted using a hemocytometer with 0.4% trypan blue staining (Corning Cat# 25-900-CI). Cells were seeded in 96-well plates at 2×10^3^ cells/well in 100 μL media and allowed to adhere for 24 h before treatment. Cells were treated with 4 μM regorafenib (Selleck Chemicals Cat# S1178), alone or in combination with 1 mM aspirin (Selleck Chemicals Cat# S3017). These concentrations are consistent with plasma regorafenib and salicylate concentrations observed clinically (29–32). Aspirin was dissolved directly in culture media, while regorafenib was dissolved in dimethyl sulfoxide (DMSO, Sigma-Aldrich Cat# 472301) then added to culture media, resulting in a final DMSO concentration of 0.1%. The total number of viable cells in each condition was estimated using the colorimetric 3-(4,5-dimethylthiazol-2-yl)-5-(3-carboxymethoxyphenyl)-2-(4-sulfo-phenyl)-2H-tetrazolium inner salt (MTS) assay. CellTiter 96 Aqueous One Solution MTS Reagent (20 μL, Promega Cat# G3582) was added to each well and incubated at 37°C for 1 h, before reading absorbance at 490 nm. Cell viability was defined as the absorbance of treated wells expressed as a percentage of the absorbance of untreated wells. Cells were also co-treated with a pan-caspase inhibitor, Z-VAD-FMK (80 μM, AdooQ Cat# A12373), to determine if the effects of aspirin and regorafenib were dependent on caspase activity. Within each experiment, triplicate wells were assayed for each treatment condition.

### Flow Cytometry

Flow cytometry experiments were performed to determine the effects of aspirin and regorafenib combination treatment on cell proliferation and cytotoxicity. Cells were seeded in T25 flasks at 1.6×10^5^ cells/flask in 8 mL media and allowed to adhere for 24 h before treatment with 1 mM aspirin and 4 μM regorafenib. Adherent cells were harvested after 48 h using TrypLE Express (ThermoFisher Cat# 12605) and pooled with non-adherent cells. Proliferation was assessed by staining for Ki67. Approximately 5×10^5^ cells were stained for 30 min using the Zombie Aqua fixable viability dye (BioLegend Cat# 423101, 1:500 dilution), then fixed for 1 h in Foxp3/Transcription Factor Fixation/Permeabilisation Solution (ThermoFisher Cat# 00552300). Cells were subsequently stained for 30 min using an APC-conjugated anti-Ki67 monoclonal antibody (ThermoFisher Cat# 14569880, RRID: AB_10854564, 1:100 dilution) in 50 μL Permeabilisation Buffer (ThermoFisher Cat# 00552300). Cells were resuspended in 2% fetal-bovine serum in PBS and analyzed using the LSRFortessa flow cytometer (BD Biosciences). A total of 20,000 events were collected for each sample. Data analysis was performed in FlowJo (v10.6.1). Debris and doublets were excluded based on the forward scatter-area over side scatter-area and side scatter-area over side scatter-height plots, respectively. The viable (Zombie Aqua-negative), Ki67-positive population was gated with reference to single-stained controls, and the median fluorescence intensity (MFI) of Ki67 staining was determined.

Cytotoxicity and apoptosis were assessed using the Annexin V-CF Blue 7-Aminoactinomycin D (7-AAD) Kit (Abcam Cat# AB214663) as per manufacturer’s protocol. A total of 20,000 events were collected for each sample. Doublets were excluded as above, and Annexin V-CF Blue and 7-AAD quadrants were gated with reference to single-stained controls. Debris were excluded based on the forward scatter-area over side scatter-area plot of the double-negative population only. Cells were determined to be non-viable if 7-AAD-positive, and early apoptotic if Annexin V-CF Blue-positive and 7-AAD-negative. Cells treated with 100 μM 5-fluorouracil were used a positive control for induction of cytotoxicity and apoptosis (Supplementary Figure S2).

### Patient-Derived Organoid Experiment

Collection of CRC patient samples for organoid derivation was approved by the Cabrini Research Governance Office (CRGO 04-19-01-15) and the Monash Human Research Ethics Committee (MHREC ID 2518). Organoids were established from treatment naïve patients diagnosed with CRC undergoing surgical resection at the Cabrini Hospital (Malvern, Australia) and cultured for experiments as previously described (33). Patients provided written informed consent. Three patient-derived organoid (PDO) lines, representing *KRAS*-mutant (G12C, PDO73T), *BRAF*-mutant (V600E, PDO118T) and *RAS/RAF*-wildtype (PDO53T) subgroups, were included in the study. After organoids were established in 24-well plates, they were dissociated using TrypLE Express and seeded as single cells in Matrigel (Corning Cat# 356231) into 96-well plates in triplicate. Organoids were cultured in complete culture medium for 4 days until small organoids were formed. Reference cell viability values were determined by adding 100 μL of Presto Blue reagent (ThermoFisher Cat# P50200) diluted in culture medium to each well. After incubating for 1 h at 37 ◦C, the fluorescence of the Presto Blue solution was measured (excitation of 560 nm and emission of 590 nm) on the PHERAstar FS (BMG Labtech). Culture medium containing 1 mM aspirin and 4 μM regorafenib, or 1 mM aspirin only, 4 μM regorafenib only or DMSO only, was replaced onto the organoids at 0 h, 24 h and 48 h. Cell viability measurements were repeated at 24 h, 48 h and 72 h. Organoids were imaged at each time point using the EVOS FL imaging system AMF4300 (ThermoFisher).

### RNA Sequencing

Total RNA was extracted from cells treated for 48 h with 1 mM aspirin and 4 μM regorafenib using the AllPrep DNA/RNA/Protein Mini Kit (Qiagen Cat# 80004) as per manufacturer’s protocol. This included on-column digestion of DNA by deoxyribonuclease I (Qiagen Cat# 79254). RNA sequencing libraries were generated using the TrueSeq Stranded mRNA Library Prep Kit (Illumina Cat# 20020594) as per manufacturer’s protocol. Libraries were sequenced across two high-output 150 cycle flow cells using the NextSeq550 instrument (Illumina), generating approximately 38M 75bp paired-end reads per sample. Sequencing data were processed using the rnaseq nf-core pipeline (pipeline v3.0, NextFlow v20.11.0). This pipeline trims reads using TrimGalore (v0.6.6), aligns reads to GRCh38 using STAR (v2.6.1d), and quantifies transcript and gene level expression using Salmon (v1.4.0). Various quality control metrics were generated by other tools including dupradar (v1.18.0), fastQC (v0.11.9), preseq (v2.0.3), qualimap (v2.2.2), rseqc (v3.0.1) and subread (v2.0.1). Gene set variation analysis (GSVA) was performed using the gsva R package (v1.34.0) to obtain sample-wise gene set enrichment scores for the Molecular Signature Database hallmark gene set collection (v7.3). All data processing was performed at the cell line level.

### Western Blotting

Whole cell protein was extracted from cells treated for 24 h with 1 mM aspirin and 4 μM regorafenib using radio-immunoprecipitation assay buffer (Cell Signaling Cat# 9806) supplemented with 1 mM phenylmethanesulfonyl fluoride (Cell Signaling Cat# 8553), 1% Protease Inhibitor Cocktail (Cell Signaling Cat# 5871) and 1% Phosphatase Inhibitor Cocktail (Cell Signaling Cat# 5870). Lysates were sonicated at 4°C using the Bioruptor Sonicator (Diagenode), with 6 cycles of 10 seconds on and 10 seconds off. The supernatant was collected after 10 min centrifugation at 16,000×g. Total protein concentration was determined using the Bicinchoninic Acid Protein Assay Kit (Sigma-Aldrich Cat# BCA1) as per manufacturer’s protocol. Western blotting was performed using the Bolt Mini Gel system (ThermoFisher). Protein samples (7.5 μg) were incubated for 10 min at 70°C with Bolt LDS Sample Buffer (ThermoFisher Cat# B0007) and Bolt Reducing Agent (ThermoFisher Cat# B0004), then loaded into 17-well 4-12% gradient Bis-Tris polyacrylamide gels (ThermoFisher Cat# NW04127). Gel electrophoresis was performed at 130 V for 70 min in Bolt MOPS SDS Running Buffer (ThermoFisher Cat# B0001). Proteins were subsequently transferred to a 0.45 μm polyvinylidene difluoride membrane, at 20 V for 90 min in Bolt Transfer Buffer (ThermoFisher Cat# BT0006) containing 10% methanol. Blocking was performed at room temperature for 1 h, using 5% bovine serum albumin in Tris-buffered saline (20 mM Tris, 150 mM NaCl, pH 7.6) containing 0.1% Tween 20 (ChemSupply Cat# TL020). Incubation with primary antibodies, diluted in 3% bovine serum albumin (Sigma-Aldrich Cat# 9647) in Tris-buffered saline with 0.1% TWEEN 20, was performed at 4°C overnight (Supplementary Table S3). Incubation with a horse radish peroxidase-conjugated anti-rabbit IgG secondary antibody (Cell Signaling Cat# #7074, RRID: AB_2099233), also diluted in 3% bovine serum albumin in Tris-buffered saline with 0.1% TWEEN 20, was performed at room temperature for 1 h. Antibody-protein complexes were visualized using the Pierce Enhanced Chemiluminescence Western Blotting Substrate (ThermoFisher Cat# 32109). Images were acquired using the ChemiDoc Imaging System (Bio-Rad). Band intensities of proteins of interest were determined using ImageJ (v2) and normalized to the corresponding β-actin or β-tubulin loading controls. Phosphorylation levels of ERK1/2 and AMPKα were quantified by expressing the normalized band intensity of the phosphorylated protein as a ratio of the normalized band intensity of the total protein.

### *In Vivo* Xenograft Growth

Animal experimentation was approved by the QIMR Berghofer Animal Ethics Committee (P302). Five million SW480 cells, suspended in Matrigel and RPMI-1640 medium at a ratio of 1:3, were injected subcutaneously into the left and right dorsal flanks of six-week-old female athymic BALB/c-Foxn1^nu^/Arc mice (Animal Resources Centre Western Australia, RRID: 2161064). Once tumors were palpable, the volumes were monitored using the standard formula, V = 0.5LW^2^, where L was the greatest length of the tumor and W was the perpendicular length measured using digital calipers. Once the average tumor volume reached approximately 200 mm^3^, the mice were randomized into four groups for treatment by daily oral gavage: vehicle only (4.5% DMSO, 30% polyethylene glycol 400 (Scharlau Cat# PO00351000) and 5% TWEEN 80 (Sigma-Aldrich Cat# P6224) in water), treatment with regorafenib only (10 mg/kg/day), treatment with aspirin only (100 mg/kg/day), or combination treatment with regorafenib (10 mg/kg/day) and aspirin (100 mg/kg/day). Tumor volume was measured every second day over 15 days of treatment. After the final day of treatment, the mice were sacrificed by cervical dislocation and the tumors were removed for histological analyses.

Tumors were fixed in formalin and embedded in paraffin. Deparaffinized and rehydrated sections were used for immunohistochemistry to detect expression of Ki67 and cleaved caspase 3. Endogenous peroxidase activity was blocked by incubating with 2% hydrogen peroxide for 10 min. For detection of Ki67, heat mediated antigen retrieval was performed for 3 min at 125°C in citrate pH 6.0 target retrieval solution (Dako Cat# S2369) using a decloaking chamber (Biocare Medical). Non-specific protein interactions were blocked by incubating with Background Sniper (Biocare Medical Cat# BS966) and 1% bovine serum albumin for 30 min. Rabbit anti-Ki67 monoclonal antibody (Abcam Cat# AB16667, RRID: AB_302459, 1:100 dilution) in Da Vinci Green diluent (Biocare Medical Cat# PD900) was applied for 30 min. For detection of cleaved caspase 3, antigen retrieval was performed for 15 min at 105°C in Tris-EDTA pH 8.8 buffer. Non-specific protein interactions were blocked by incubating with 10% goat serum (Sigma-Aldrich Cat# S26) for 30 min. Rabbit anti-cleaved caspase 3 monoclonal antibody (Biocare Medical Cat# CP229, RRID:AB_2737391, 1:100 dilution) was applied for 2 h. The MACH 1 Universal HRP-Polymer Detection Kit (Biocare Medical Cat# M1U539) was used for signal development. After hematoxylin counterstaining and mounting, the sections were scanned at 40X magnification using the Aperio AT Turbo bright-field slide scanner (Leica). The percentages of viable tumor cells positive for Ki67 and cleaved caspase 3 were quantified using QuPath (v0.2.3).

### Statistical Analysis

Statistical analysis was performed using JMP Pro (v15.2.0). All data were log-transformed prior to statistical analysis, with exception of the gene set enrichment scores determined from RNA sequencing. The two-way ANOVA was used to test for interaction between aspirin treatment and regorafenib treatment. Tukey’s post-hoc test was performed to compare the aspirin and regorafenib combination treatment condition to the untreated control, aspirin single treatment and regorafenib single treatment conditions. Additionally, interaction p values from analysis of the RNA sequencing data were adjusted using the Benjamini-Hochberg method to control for false discovery rate. For experiments involving data collection at more than one time point, data from the final time point were used for statistical analysis. All plots were generated using GraphPad Prism (v8.2.1).

## Results

### Aspirin and regorafenib synergistically inhibit proliferation of colorectal cancer cell lines *in vitro*

RKO, SW480 and LIM1215 CRC cells were treated with 1 mM aspirin and 4 μM regorafenib to compare the effects of combination treatment to either treatment alone. Cell viability was measured using the MTS assay after 24, 48 and 72 h, and normalized to the untreated control at each time point. The two-way ANOVA showed significant interactions between the effects of aspirin treatment and regorafenib treatment on cell viability after 72 h (interaction p < 0.01 for all cell lines; Figure 1A–1C). Combination treatment with 1 mM aspirin and 4 μM regorafenib significantly reduced cell viability compared to the untreated control, to a greater extent than single treatment with either 1 mM aspirin (p < 0.0001) or 4 μM regorafenib (p < 0.0001 for all cell lines; Figure 1A–1C). These results indicate that aspirin and regorafenib interact synergistically to reduce CRC cell viability. This is consistent with an initial screen of a greater range of aspirin and regorafenib concentrations and a greater number of cell lines, which had suggested that addition of aspirin, at concentrations consistent with plasma salicylate concentrations observed clinically (250 μM to 2 mM) (29–32), could enhance the effects of regorafenib on CRC cells (Supplementary Figure S4).

**Figure 1:**
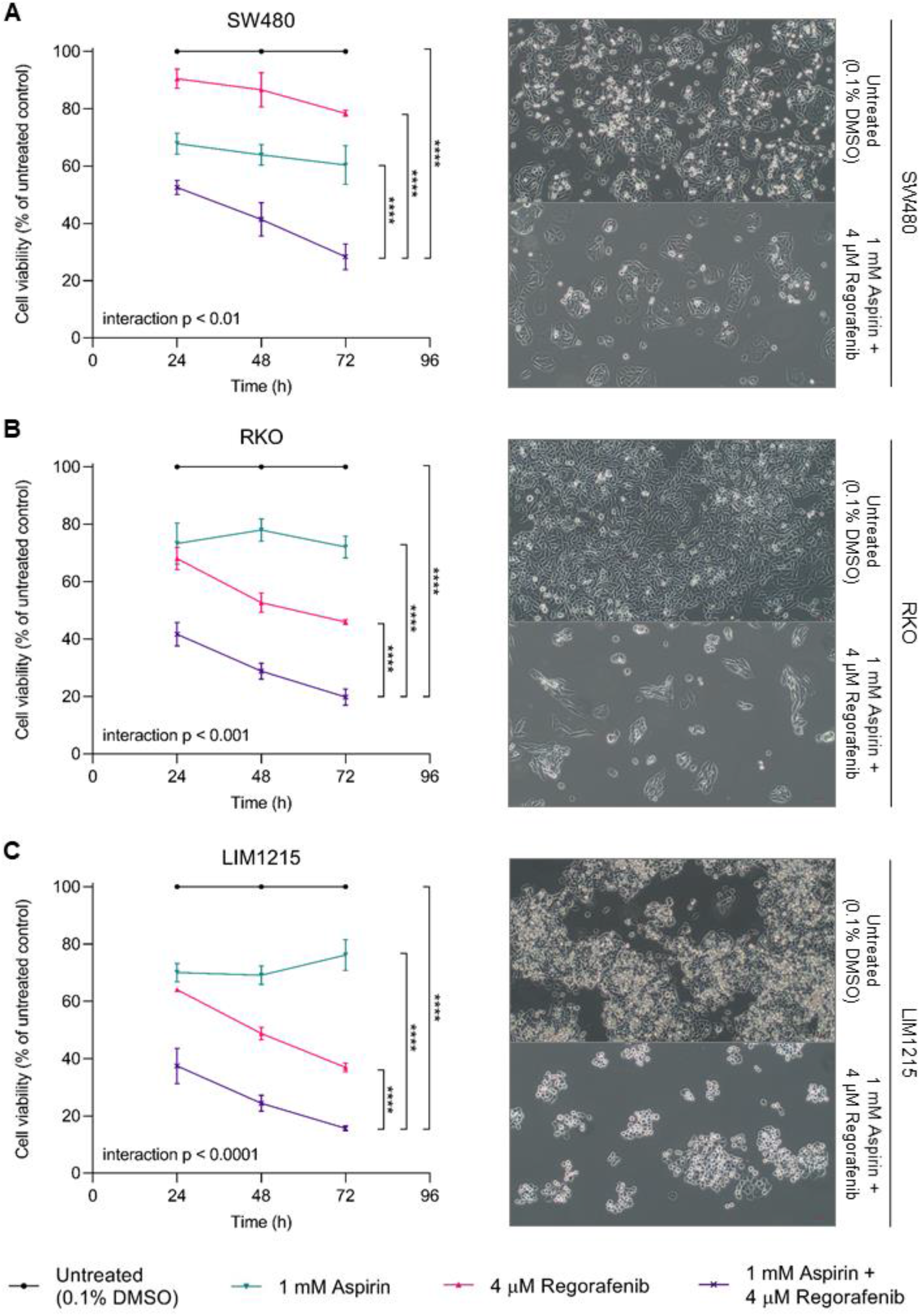
Aspirin and regorafenib synergistically reduce viability of colorectal cancer cells *in vitro.* Cell viability was measured after 24 h, 48 h and 72 h treatment with 1 mM aspirin and/or 4 μM regorafenib using the MTS assay and normalized to the untreated control. Data represent mean ± SD (n = 3). Statistical analysis was performed at the 72 h time point using the two-way ANOVA and Tukey’s post-hoc test following log transformation (****, p < 0.0001). Representative images (10X objective) taken at the 72 h time point are shown for the combination treatment and untreated control conditions. **(A)** SW480 (*KRAS*-mutant). **(B)** RKO (*BRAF*-mutant). **(C)** LIM1215 (*RAS/RAF*-wildtype).

Proliferation was assessed by Ki67 staining and flow cytometry. The distribution of Ki67 expression levels within the viable, Ki67-positive population was altered after 48 hours of treatment with 1 mM aspirin and 4 μM regorafenib (Figure 2A). There were significant interactions between the effects of aspirin and regorafenib on the median fluorescence intensity (MFI) of Ki67 staining (interaction p < 0.01 for all cell lines; Figure 2B). Ki67 MFI of each cell line was significantly lower following aspirin and regorafenib combination treatment, compared to the untreated control (p < 0.0001) and compared to single treatment with either aspirin (p < 0.0001) or regorafenib (p < 0.001 for all cell lines; Figure 2B). This demonstrates that aspirin and regorafenib synergistically inhibit proliferation of CRC cells.

**Figure 2:**
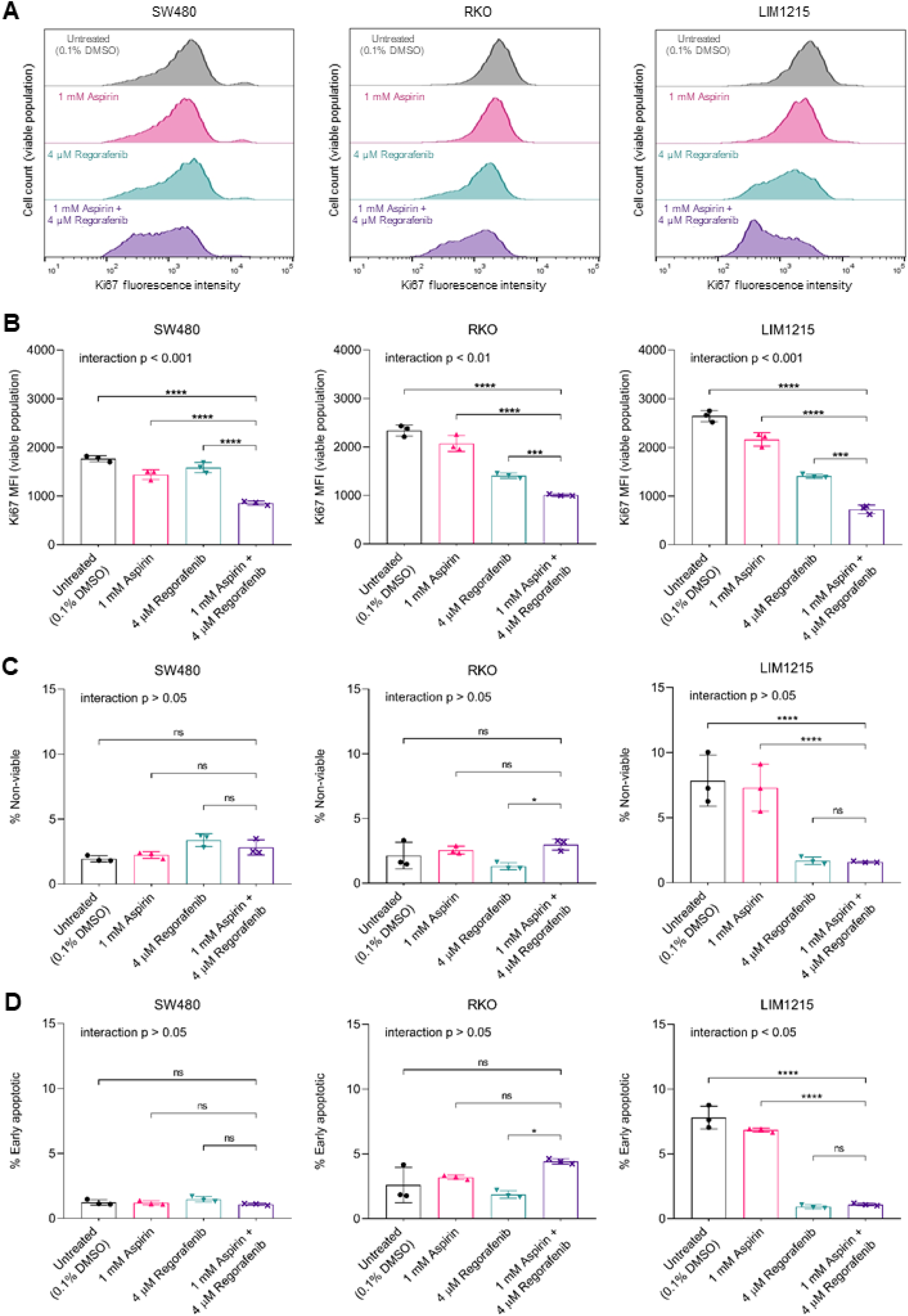
Aspirin and regorafenib synergistically inhibit proliferation of colorectal cancer cells *in vitro* without inducing cytotoxicity. Flow cytometry was performed after 48 h treatment with 1 mM aspirin and/or 4 μM regorafenib. Data represent mean ± SD (n = 3). Statistical analysis was performed using the two-way ANOVA and Tukey’s post-hoc test following log transformation (ns, p > 0.05; *, p < 0.05; **, p < 0.01; ***, p <0.001; ****, p < 0.0001). **(A)** Ki67 median fluorescence intensity of viable cells. **(B)** Representative distributions of Ki67 expression. **(C)** Percentage of non-viable (7-AAD-positive) cells. **(D)** Percentage of early apoptotic (Annexin V-positive and 7-AAD-negative) cells.

Cytotoxicity and apoptosis were assessed by flow cytometry. There were no differences in the percentages of non-viable (7-AAD-positive) and early apoptotic (Annexin V-positive and 7-AAD-negative) SW480 cells across the treatment conditions after 48 h (Figure 2C–2D). The percentages of non-viable and early apoptotic RKO cells were greater after combination treatment compared to regorafenib treatment alone (p < 0.05), but not different to the untreated control or aspirin single treatment (Figure 2C–2D). Additionally, the percentages of non-viable and early apoptotic LIM1215 cells were decreased after combination treatment compared to the untreated control (p < 0.01), however there were no differences between the combination treatment and regorafenib single treatment (Figure 2C–2D). These results suggest that combination treatment of CRC cell lines with 1 mM aspirin and 4 μM regorafenib does not induce significant levels of cytotoxicity or apoptosis *in vitro*. Additionally, co-treatment with the pan-caspase inhibitor Z-VAD-FMK (80 μM) did not rescue cell viability (Supplementary Figure S5), showing that the effects of combination treatment at these concentrations *in vitro* are caspase-independent, consistent with the lack of apoptosis observed. Therefore, the synergistic reduction in cell viability observed using the MTS assay *in vitro* can be attributed to synergistic inhibition of proliferation rather than induction of cytotoxicity or apoptosis.

### Aspirin and regorafenib synergistically inhibit proliferation of patient-derived colorectal cancer organoids

Aspirin and regorafenib combination treatment was also tested in three-dimensional patient-derived organoid models. Combination treatment with 1 mM aspirin and 4 μM regorafenib reduced the viability of three different patient-derived organoid lines after 72 h (p < 0.0001 for all organoid lines; Figure 3A–3C). Greater reductions in viability were observed following combination treatment, compared to single treatment with either 1 mM aspirin (p < 0.0001) or 4 μM regorafenib (p < 0.001 for all organoid lines; Figure 3A–3C). These differences were independent of MAPK pathway mutation status, as the three CRC samples from which the organoids were derived represented *KRAS*-mutant (G12C), *BRAF*-mutant (V600E) and *RAS/RAF*-wildtype subgroups. There were significant interactions between the effects of aspirin and regorafenib (interaction p < 0.01 for all organoid lines; Figure 3A–3C), showing that the synergy observed in CRC cell lines extends to three-dimensional patient-derived CRC organoids.

**Figure 3:**
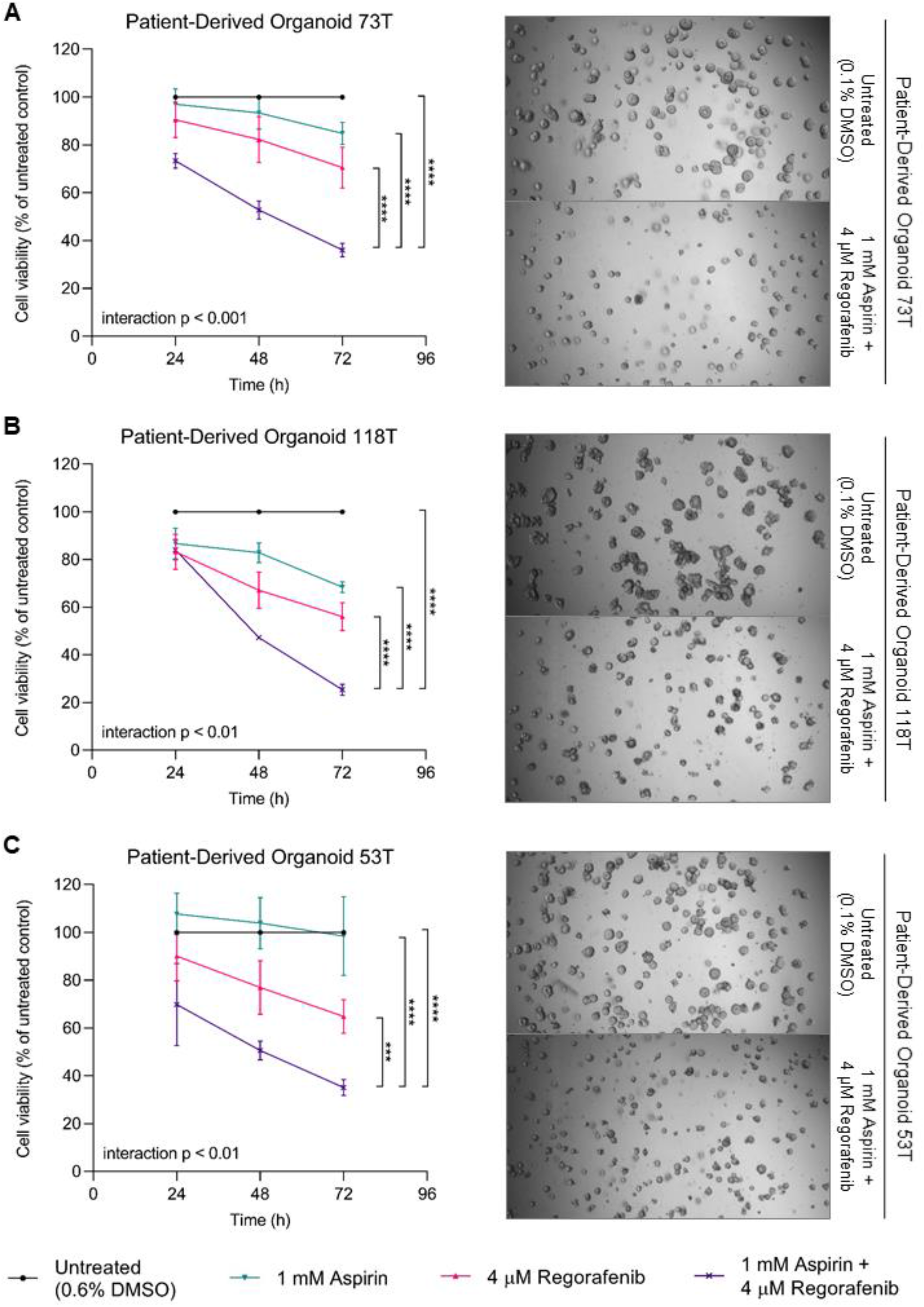
Aspirin and regorafenib synergistically reduce viability of patient-derived colorectal cancer organoids. Cell viability was measured after 24 h, 48 h and 72 h treatment with 1 mM aspirin and/or 4 μM regorafenib using the Presto Blue assay and normalized to the untreated control. Data represent mean ± SD (n = 3). Statistical analysis was performed at the 72 h time point using the two-way ANOVA and Tukey’s post-hoc test following log transformation ***, p < 0.001; ****, p < 0.0001). Representative images (4X objective) taken at the 72 h time point are shown for the combination treatment and untreated control conditions. **(A)** Patient-derived organoid 73T (*KRAS*-mutant). **(B)** Patient-derived organoid 118T (*BRAF*-mutant). **(C)** Patient-derived organoid 53T (*RAS/RAF*-wildtype).

### Aspirin and regorafenib combination treatment alters molecular pathways within colorectal cancer cells

RNA sequencing was performed to explore the molecular effects of aspirin and regorafenib combination treatment on CRC cells (Figure 4A–4B). The SW480 and RKO cell lines showed minimal expression of cyclooxygenase 2 (COX2), which is the target of aspirin that mediates its anti-inflammatory and analgesic effects. On RNA sequencing, *COX2* mRNA was undetectable in SW480 cells and 0.07 transcripts per million in RKO cells. As SW480 and RKO are sensitive to aspirin and regorafenib treatment despite minimal COX2 expression, inhibition of COX2 is unlikely to be critical for the synergistic anti-proliferative effect.

**Figure 4:**
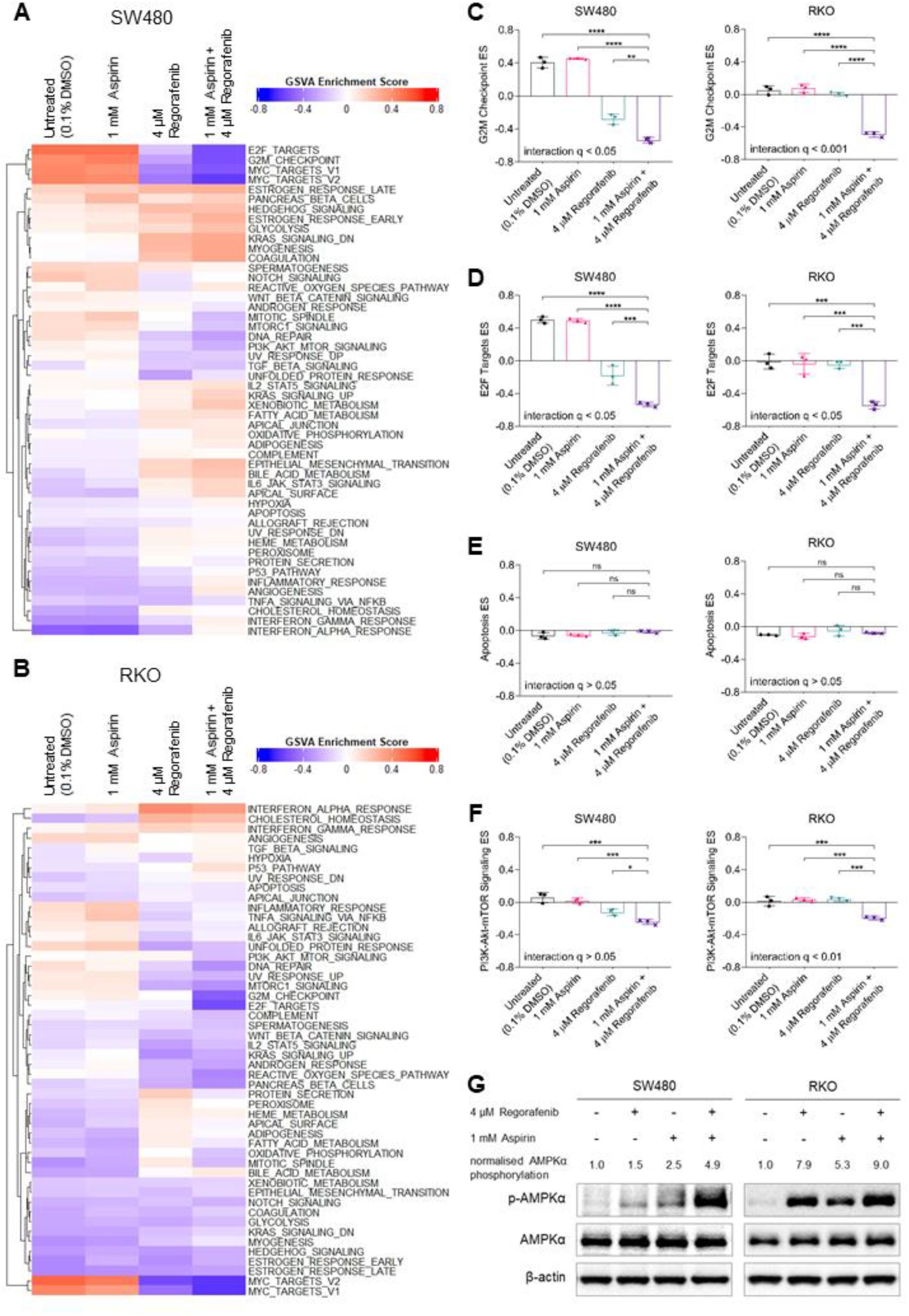
Aspirin and regorafenib combination treatment induces molecular changes in colorectal cancer cells *in vitro.* RNA sequencing was performed after 48 h treatment with 1 mM aspirin and/or 4 μM regorafenib and Western blotting was performed after 24 h treatment. **(A)** Gene set enrichment analysis for gene sets in the Molecular Signature Database hallmark collection. **(B-F)** Enrichment scores for “G2M Checkpoint”, “E2F Targets”, “Apoptosis”, and “PI3K-Akt-mTOR Signaling” gene sets. Data represent mean ± SD (n = 3). Statistical analysis was performed using the two-way ANOVA and Tukey’s post-hoc test(ns, p > 0.05; *, p < 0.05; **, p < 0.01; ***, p < 0.001; ****, p < 0.0001). Interaction p values were adjusted using the Benjamini-Hochberg method to correct for false discovery rate (q). **(G)** AMPK activation (ratio of phosphorylated AMPKα to total AMPKα).

Gene set enrichment analysis showed that combination treatment with aspirin and regorafenib induced downregulation of the “G2M Checkpoint” and “E2F Targets” gene sets from the Molecular Signature Database hallmark gene set collection (p < 0.001), with significant interactions between aspirin treatment and regorafenib treatment after adjustment for false discovery rate (interaction q < 0.05 for both hallmarks and cell lines; Supplementary Table S6; Figure 4C–4D). These hallmarks were downregulated to a greater extent following combination treatment compared to treatment with aspirin (p < 0.001) or regorafenib alone (p < 0.01 for both hallmarks and cell lines; Figure 4C–4D), consistent with the synergistic anti-proliferative effect observed. Additionally, there were no differences in enrichment scores for the “Apoptosis” hallmark across the treatment conditions for SW480 or RKO (Figure 4E). These results support the inhibition of proliferation and lack of apoptosis observed by flow cytometry *in vitro*.

The effects of aspirin and regorafenib combination treatment on enrichment scores for the “KRAS Signaling” hallmarks were variable between the *KRAS*-mutant SW480 cells and *BRAF*-mutant RKO cells, with no significant interactions, as were the effects on ERK1/2 phosphorylation levels determined by Western blotting (Supplementary Figure S7). In contrast, the “PI3K-Akt-mTOR signaling” hallmark was consistently downregulated in SW480 and RKO cells following aspirin and regorafenib combination treatment (p < 0.001 for both cell lines; Figure 4F). There was a significant interaction between the effects of aspirin treatment and regorafenib treatment on the hallmark in *PIK3CA*-mutant RKO cells (q < 0.05) but not in *PIK3CA*-wildtype SW480 cells (Figure 4F). However, downregulation was observed in both cell lines to a greater extent following combination treatment than following single treatment with either aspirin (p < 0.001) or regorafenib (p < 0.05 for both cell lines; Figure 4F). Western blotting for phosphorylated AMPKα showed that these changes were also associated with increased AMPK activation (Figure 4G). These pathways may contribute to the synergistic interaction between aspirin and regorafenib.

### Aspirin and regorafenib combination treatment effectively inhibits growth of SW480-derived tumors *in vivo*

Aspirin and regorafenib combination treatment synergistically inhibited growth of SW480-derived tumors in athymic mice (interaction p < 0.05; Figure 5A). Tumors from two mice were excluded from analyses due to early sacrifice following procedure-related adverse events. Tumor volumes were lower following 15 days of aspirin (100 mg/kg/day) and regorafenib (10 mg/kg/day) combination treatment compared to 15 days of aspirin (p < 0.0001) or regorafenib treatment alone (p < 0.001), or vehicle alone (p < 0.0001; Figure 5A). A greater percentage of viable cells within tumors from the combination treatment condition were positive for cleaved caspase 3, compared to tumors from the untreated control condition (p < 0.0001) and aspirin single treatment condition (p < 0.0001; Figure 5B). This shows that aspirin and regorafenib combination treatment induces apoptosis of CRC cells *in vivo*. However, there was no difference between the combination treatment and regorafenib single treatment conditions, consistent with the lack of interaction effect observed (Figure 5B), indicating that the pro-apoptotic effect of combination treatment *in vivo* is driven by the effect of regorafenib. Additionally, combination treatment with aspirin and regorafenib reduced the percentage of viable cells that were positive for Ki67, compared to the untreated control (p < 0.001) and compared to aspirin (p < 0.0001) or regorafenib treatment alone (p < 0.05), with a significant interaction effect (interaction p < 0.05; Figure 5C). This indicates that aspirin and regorafenib synergistically inhibit proliferation of SW480 cells *in vivo*, contributing to the observed reduction in tumor volume.

**Figure 5:**
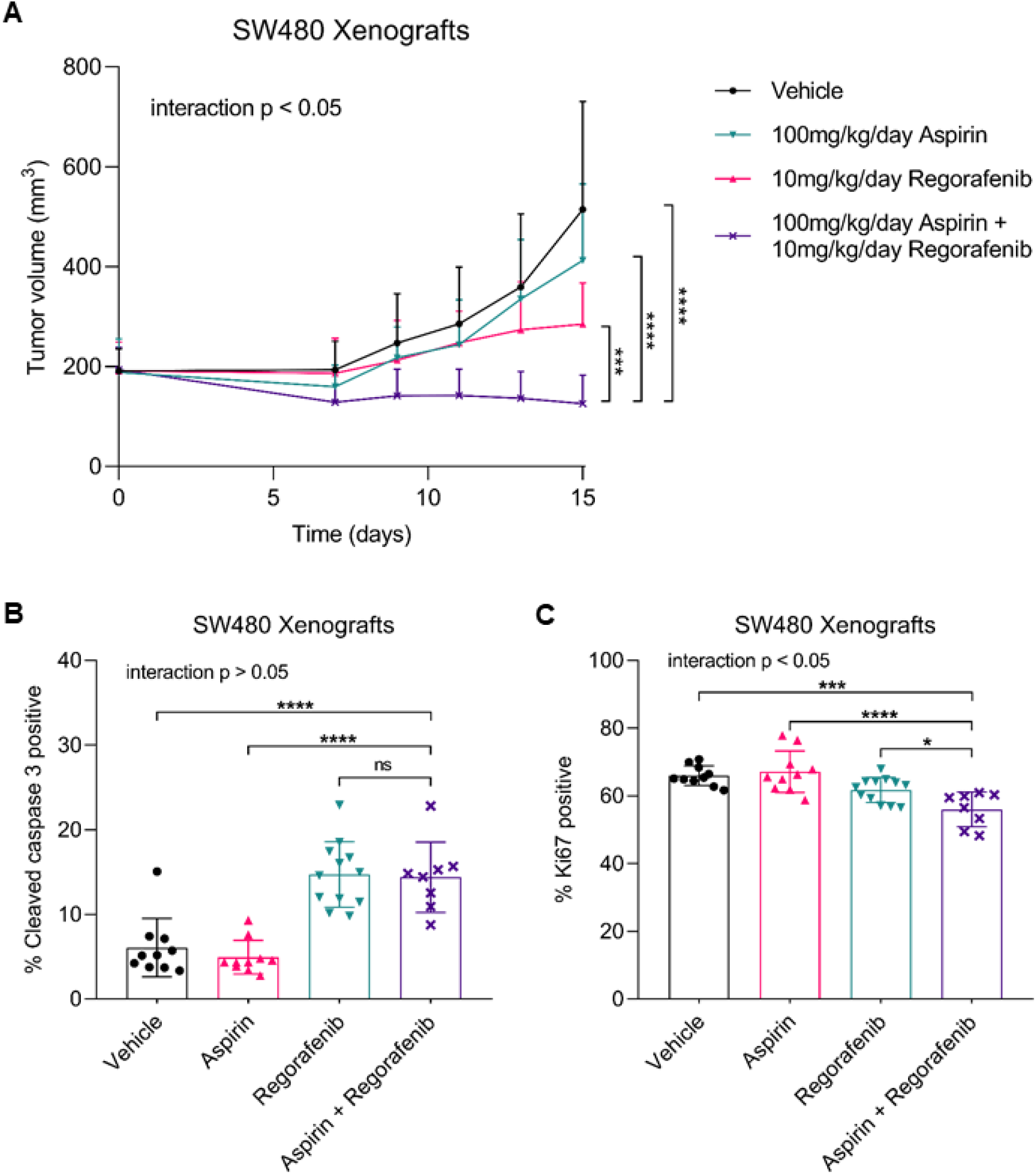
Aspirin and regorafenib synergistically inhibit growth of SW480-derived tumors *in vivo.* Xenografts were established by injecting 5×10^6^ SW480 cells into dorsal flanks of athymic BALB/c-Foxn1^nu^ mice. Mice were treated by daily oral gavage with 100 mg/kg/day aspirin and/or 10 mg/kg/day regorafenib for 15 days. Data represent mean ± SD (n = 8 to 12 per condition). Statistical analysis was performed at the experiment endpoint using the two-way ANOVA and Tukey’s post-hoc test following log transformation (ns, p > 0.05; *, p < 0.05; ***, p < 0.001; ****, p < 0.0001). **(A)** Tumor volumes over time. **(B-C)** Percentages of viable tumor cells positive for cleaved caspase 3 or Ki67 on immunohistochemistry.

## Discussion

Regorafenib is approved as a last-line treatment option for patients with metastatic CRC. However, as a monotherapy, regorafenib offers only a small improvement in overall survival with substantial toxicity (10–12). Our study explored the possibility of combining regorafenib with aspirin as a novel treatment strategy. Aspirin and regorafenib have individually been shown to inhibit proliferation and induce apoptosis of CRC cells, via complex mechanisms that remain incompletely understood (4–9, 34–37). While we and others have found the combination of aspirin and the regorafenib analogue, sorafenib, to be effective in preclinical studies of CRC, melanoma, lung carcinoma and hepatocellular carcinoma (19–23), no data for the combination of aspirin and regorafenib have previously been published. Our results demonstrate for the first time a synergistic interaction between aspirin and regorafenib in inhibiting proliferation of CRC cells *in vitro* and *in vivo*.

The synergistic anti-proliferative effect of aspirin and regorafenib was observed across cell lines representing *KRAS*-mutant, *BRAF*-mutant and *RAS/RAF-*wildtype subgroups of CRC. This is consistent with previous studies that have found no correlation between *KRAS* or *BRAF* mutation status of CRC cell lines and their sensitivity to aspirin and sorafenib combination treatment (19, 20). Additionally, RNA sequencing showed downregulation of proliferation-associated gene sets following combination treatment with aspirin and regorafenib. The synergistic anti-proliferative effect was also observed in three-dimensional patient-derived CRC organoids, which have been shown to recapitulate tumor heterogeneity and patient responses to treatment (38, 39). Therefore, combining regorafenib with aspirin may achieve improved anti-tumor activity against CRC. This is consistent with observational studies conducted by Giampieri *et al.* (17) and Casadei-Gardini *et al.* (18), which identified associations between incidental aspirin use and improved overall survival in patients with refractory metastatic CRC or hepatocellular carcinoma receiving treatment with regorafenib.

As the mechanisms underlying the effects of aspirin and regorafenib on CRC cells are complex and not well understood, we performed RNA sequencing to identify pathways that may be involved in the response to combination treatment. Inhibition of cyclooxygenase-2 (COX2) mediates aspirin’s anti-inflammatory and analgesic effects (40). However, COX2 is minimally expressed in SW480 or RKO cells (25), despite sensitivity to aspirin and regorafenib combination treatment. Therefore, COX2 inhibition is unlikely to mediate the synergistic anti-proliferative effect. This is consistent with our previous finding that COX1 and COX2 inhibitors other than aspirin, including ibuprofen and celecoxib, do not interact synergistically with sorafenib in melanoma cell lines (21). Additionally, aspirin and regorafenib combination treatment was observed to elicit variable effects on MAPK pathway activity. Variable effects on MAPK pathway activity have also been reported for the combination of aspirin and sorafenib (20, 21). While regorafenib and sorafenib inhibit the MAPK pathway in *BRAF*-mutant cells (8), paradoxical MAPK pathway activation has previously been reported in *BRAF*-wildtype cells, where binding of the inhibitor to Raf proteins promotes activation by Ras (41).

In contrast, downregulation of the PI3K-Akt-mTOR pathway was consistently observed in CRC cells following aspirin and regorafenib combination treatment, to a greater extent than either treatment alone. Synergistic interaction between the effects of aspirin and regorafenib on this pathway was observed in *PIK3CA*-mutant RKO cells but not in *PIK3CA*-wildtype SW480 cells. Enhanced AMPK activity was also consistently observed. Previous studies have demonstrated a role for PI3K-Akt-mTOR pathway inhibition and AMPK pathway activation in the anti-tumor effects of aspirin (35, 37) and regorafenib individually (22, 42). Both pathways have also been implicated in the synergistic effects of aspirin and sorafenib (20–22). Activating *PIK3CA* mutations have been identified as a marker of increased sensitivity of CRC cells to aspirin treatment *in vitro* (34, 37), and increased association between aspirin use and improved CRC survival outcomes in observational studies (16). Additionally, high levels of tissue AMPK activation were found to be associated with improved overall survival in a cohort of metastatic CRC patients treated with combination chemotherapy and bevacizumab (43). Our results align with these findings and suggest that PI3K-Akt-mTOR pathway inhibition and AMPK pathway activation may contribute to the synergistic anti-proliferative effect of aspirin and regorafenib on CRC cells.

While our *in vitro* results highlight a role for aspirin and regorafenib combination therapy in CRC irrespective of *KRAS* or *BRAF* mutation status, the *KRAS*-mutant subtype is of particular interest given the limited treatment options currently available. Activating *RAS* mutations, which are present in approximately 50% of CRCs, are predictive of resistance to anti-epidermal growth factor receptor (EGFR) therapy (44). Combination therapy with cetuximab (anti-EGFR monoclonal antibody) and encorafenib (B-Raf inhibitor) has recently emerged as an additional line of therapy for patients with metastatic *BRAF* V600E-mutant CRC (45). However, development of targeted therapy for *KRAS*-mutant CRC remains a challenge (46). Additionally, most CRCs are microsatellite stable, which is predictive of poor response to current immunotherapy approaches (47). Our results show that combination treatment with aspirin and regorafenib effectively inhibits growth of microsatellite stable *KRAS*-mutant SW480-derived tumors in an *in vivo* xenograft model. As regorafenib monotherapy is currently approved as a last-line treatment option for refractory metastatic CRC (3), this further suggests that combining regorafenib with aspirin may be an improved treatment strategy.

Overall, aspirin and regorafenib demonstrate synergistic anti-proliferative effects in preclinical models of CRC. Synergistic inhibition of the PI3K-Akt-mTOR pathway and activation of the AMPK pathway may mediate these effects. Further molecular studies are required for better characterization of the underlying mechanisms. These findings provide a rationale for investigating aspirin and regorafenib combination therapy as a potential treatment strategy for patients with refractory metastatic CRC.

