## Supplementary Tables and Figures for "Aspirin Synergizes with Regorafenib to Reduce Growth of Colorectal Cancer"

**Supplementary Table S1: Molecular characteristics of RKO, SW480 and LIM1215 colorectal cancer cell lines.**

| Cell Line | Site | *KRAS* [a] | *BRAF* [a] | *TP53* [a] | *PIK3CA* [a] | MSI [b] | CIMP [b] |
|---|---|---|---|---|---|---|---|
| **RKO** | Primary tumor | wt | V600E | wt | H1047R | Unstable | Positive |
| **SW480** | Primary tumor | G12V | wt | R273H, P309S | wt | Stable | Negative |
| **LIM1215** | Omental metastasis | wt | wt | wt | wt | Unstable | Negative |

[a] Mutation data for *KRAS, BRAF, TP53* and *PIK3CA* were obtained from the Broad Institute Cancer Cell Line Encyclopedia for RKO and SW480 (25), with changes depicted at the amino acid level (wt, wild-type). Mutation data for LIM1215 were reported by Zhang *et al.* (26).

[b] Microsatellite instability (MSI) and CpG island methylator phenotype (CIMP) data for RKO and SW480 were obtained from a comprehensive study of 34 colorectal cancer cell lines conducted by Berg *et al*. (27). MSI and CIMP data for LIM1215 were reported by Zhang *et al.* (26) and Suter *et al.* (28) respectively.

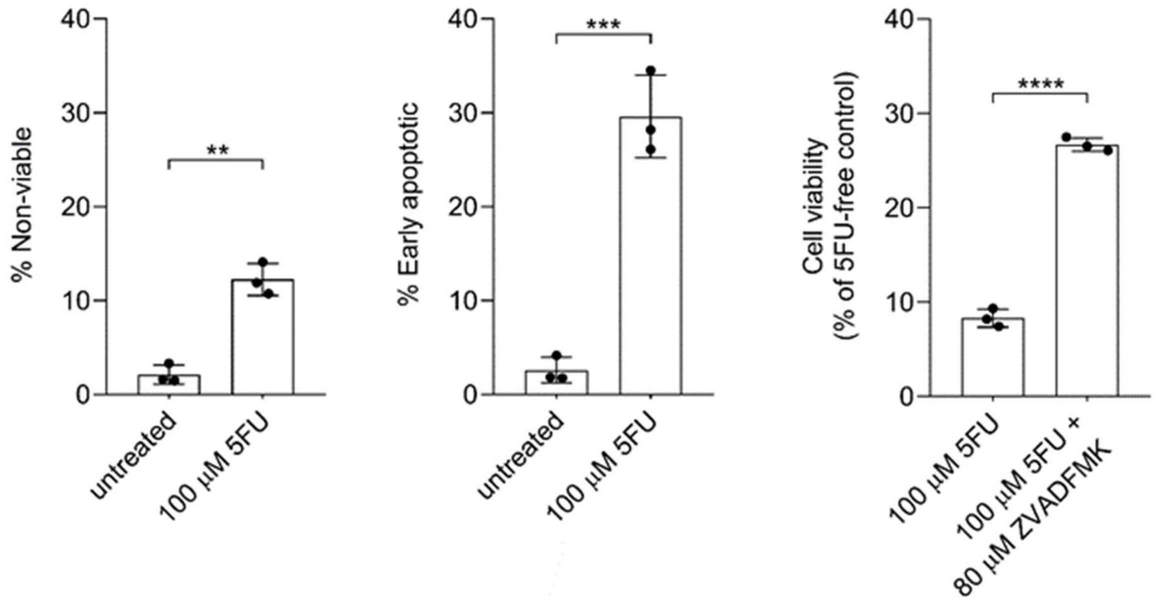


**Supplementary Figure S2: RKO colorectal cancer cells treated with 100 µM 5-fluorouracil as positive controls for cytotoxicity and apoptosis.** Percentages of non-viable (7-AAD-positive) and early apoptotic (Annexin V-positive and 7-AAD-negative) cells after 48 h treatment were determined using flow cytometry. Cell viability after 72 h co-treatment with 80 µM pan-caspase inhibitor Z-VAD-FMK was measured using the MTS assay. Data represent mean ± SD (n=3). Statistical analysis was performed using the unpaired t-test (**, $p < 0.01$; ***, $p < 0.001$; ****, $p < 0.0001$).

**Supplementary Table S3: Primary antibodies used for Western blotting to detect protein expression and phosphorylation.**

| Target Protein [a] | Antibody Type | Supplier, Catalogue Number, RRID | Dilution Factor |
|---|---|---|---|
| **AMPKα** | Rabbit IgG monoclonal | Cell Signaling Cat# 5831, RRID: AB_10622186 | 1:1000 |
| **Phospho-AMPKα (Thr172)** | Rabbit IgG monoclonal | Cell Signaling Cat# 2535, RRID: AB_331250 | 1:1000 |
| **ERK1/2** | Rabbit IgG monoclonal | Cell Signaling Cat# 4695, RRID: AB_390779 | 1:1000 |
| **Phospho-ERK1/2 (Thr202/Tyr204)** | Rabbit IgG monoclonal | Cell Signaling Cat# 4370, RRID: AB_2315112 | 1:2000 |
| **β-actin (loading control)** | Rabbit IgG polyclonal | Cell Signaling Cat# 4967, RRID: AB_330288 | 1:1000 |
| **β-tubulin (loading control)** | Rabbit IgG polyclonal | Abcam Cat# 6046, RRID: AB_2210370 | 1:2000 |

[a] Antibodies were used to detect total and phosphorylated AMPK (5’-adenosine monophosphate-activated protein kinase) and ERK (extracellular signal-regulated kinase) levels.

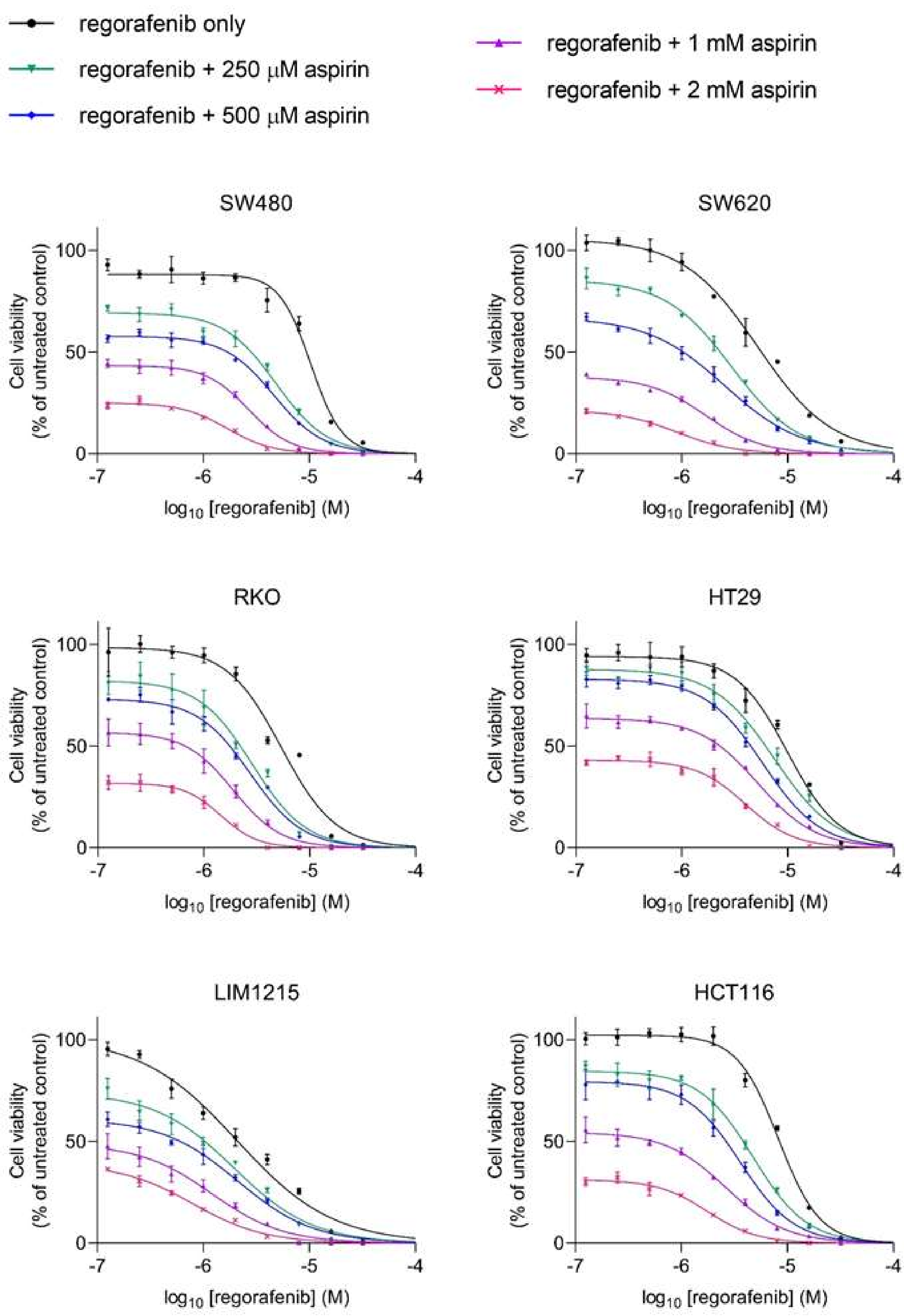


**Supplementary Figure S4: Concentration-effect curves for regorafenib (125 nM to 32 µM) in the presence of physiological concentrations of aspirin (250 µM to 2 mM), for six colorectal cancer cell lines.** Cell viability was measured after 72 h treatment using the MTS assay and normalized to the untreated control. Data represent mean ± SD of triplicate wells.

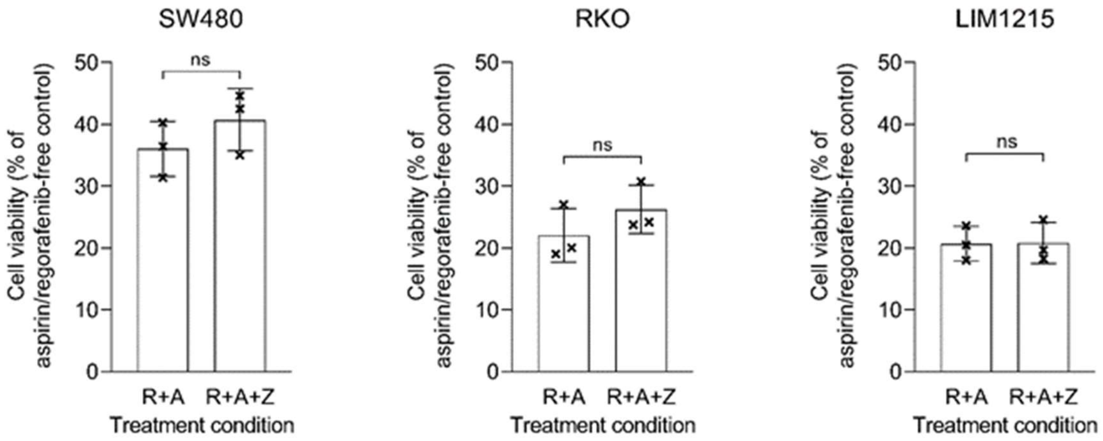


**Supplementary Figure S5: Colorectal cancer cells treated with 1 mM aspirin and 4 µM regorafenib (R+A) and co-treated with 80 µM pan-caspase inhibitor Z-VAD-FMK (R+A+Z).** Cell viability was measured after 72 h treatment using the MTS assay. Data represent mean ± SD (n=3). Statistical analysis was performed using the unpaired t-test (ns, $p > 0.05$).

**Supplementary Table S6: Gene set variation analysis of colorectal cancer cells following treatment with 1 mM aspirin and/or 4 µM regorafenib.** RNA sequencing and gene set variation analysis using the Molecular Signature Database hallmark collection was performed after 48 h treatment (n=3). Statistical analysis was performed using the two-way ANOVA for aspirin treatment and regorafenib treatment. Interaction p values were adjusted using the Benjamini-Hochberg method to correct for false discovery rate (italicized if q < 0.05).

| Hallmark Gene Set | Adjusted interaction p value (q) | |
|---|---|---|
| | SW480 | RKO |
| HALLMARK_ADIPOGENESIS | 0.542593 | 0.881707 |
| HALLMARK_ALLOGRAFT_REJECTION | 0.175000 | 0.887500 |
| HALLMARK_ANDROGEN_RESPONSE | 0.308824 | 0.422917 |
| HALLMARK_ANGIOGENESIS | 0.210000 | 0.894318 |
| HALLMARK_APICAL_JUNCTION | 0.827500 | 0.434783 |
| HALLMARK_APICAL_SURFACE | 0.550000 | 0.176154 |
| HALLMARK_APOPTOSIS | 0.959574 | 0.925000 |
| HALLMARK_BILE_ACID_METABOLISM | 0.960870 | 0.051611 |
| HALLMARK_CHOLESTEROL_HOMEOSTASIS | 0.610000 | 0.488889 |
| HALLMARK_COAGULATION | 0.189643 | 0.817143 |
| HALLMARK_COMPLEMENT | 0.828205 | 0.056786 |
| HALLMARK_DNA_REPAIR | 0.381579 | *0.004338* |
| HALLMARK_E2F_TARGETS | *0.029333* | *0.013200* |
| HALLMARK_EPITHELIAL_MESENCHYMAL_TRANSITION | 0.390476 | 0.619355 |
| HALLMARK_ESTROGEN_RESPONSE_EARLY | 0.538462 | 0.880000 |
| HALLMARK_ESTROGEN_RESPONSE_LATE | 0.929592 | 0.974000 |
| HALLMARK_FATTY_ACID_METABOLISM | 0.532143 | 0.234063 |
| HALLMARK_G2M_CHECKPOINT | *0.034600* | *0.000248* |
| HALLMARK_GLYCOLYSIS | 0.616176 | 0.640625 |
| HALLMARK_HEDGEHOG_SIGNALING | 0.693056 | 0.904651 |
| HALLMARK_HEME_METABOLISM | 0.617188 | 0.238333 |
| HALLMARK_HYPOXIA | 0.463043 | 0.068636 |
| HALLMARK_IL2_STAT5_SIGNALING | 0.721053 | 0.872368 |
| HALLMARK_IL6_JAK_STAT3_SIGNALING | 0.943750 | 0.595000 |
| HALLMARK_INFLAMMATORY_RESPONSE | *0.039250* | 0.957143 |
| HALLMARK_INTERFERON_ALPHA_RESPONSE | *0.046417* | 0.289412 |
| HALLMARK_INTERFERON_GAMMA_RESPONSE | 0.091429 | 0.074000 |
| HALLMARK_KRAS_SIGNALING_DN | 0.918182 | 0.362500 |
| HALLMARK_KRAS_SIGNALING_UP | 0.131500 | 0.735294 |
| HALLMARK_MITOTIC_SPINDLE | 0.147727 | *0.001555* |
| HALLMARK_MTORC1_SIGNALING | 0.122222 | 0.302778 |
| HALLMARK_MYC_TARGETS_V1 | 0.162308 | 0.354762 |
| HALLMARK_MYC_TARGETS_V2 | *0.049500* | 0.885897 |
| HALLMARK_MYOGENESIS | 0.536000 | 0.347727 |
| HALLMARK_NOTCH_SIGNALING | 0.612121 | 0.870270 |
| HALLMARK_OXIDATIVE_PHOSPHORYLATION | 0.108125 | 0.494643 |
| HALLMARK_P53_PATHWAY | 0.400000 | 0.057500 |
| HALLMARK_PANCREAS_BETA_CELLS | 0.693243 | 0.240357 |
| HALLMARK_PEROXISOME | 0.520690 | 0.100833 |
| HALLMARK_PI3K_AKT_MTOR_SIGNALING | 0.405000 | *0.005000* |
| HALLMARK_PROTEIN_SECRETION | 0.854762 | 0.051938 |
| HALLMARK_REACTIVE_OXYGEN_SPECIES_PATHWAY | 0.832927 | 0.357895 |
| HALLMARK_SPERMATOGENESIS | 0.843023 | 0.631818 |
| HALLMARK_TGF_BETA_SIGNALING | 0.543333 | 0.463462 |
| HALLMARK_TNFA_SIGNALING_VIA_NFKB | 0.246250 | 0.915556 |
| HALLMARK_UNFOLDED_PROTEIN_RESPONSE | *0.022875* | 0.500000 |
| HALLMARK_UV_RESPONSE_DN | 0.413636 | 0.910638 |
| HALLMARK_UV_RESPONSE_UP | 0.937000 | 0.478000 |
| HALLMARK_WNT_BETA_CATENIN_SIGNALING | 0.902222 | 0.922826 |
| HALLMARK_XENOBIOTIC_METABOLISM | 0.545833 | 0.957292 |

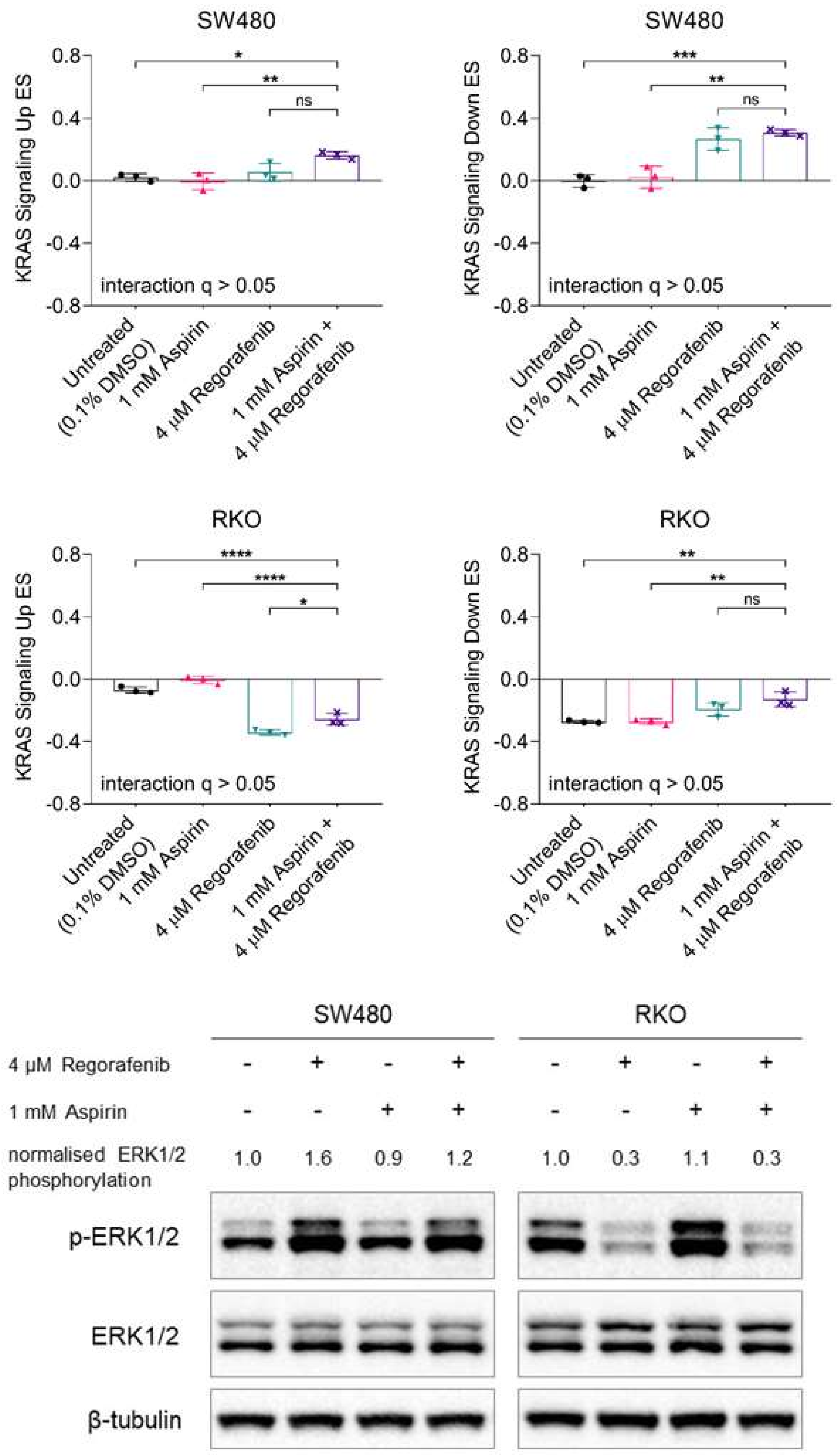


**Supplementary Figure S7: Changes in MAPK signaling in colorectal cancer cells following treatment with 1 mM aspirin and/or 4 µM regorafenib.** RNA sequencing and gene set variation analysis for the "KRAS Signaling" gene sets in the Molecular Signature Database hallmark collection was performed after 48 h treatment. Data represent mean ± SD (n=3). Statistical analysis was performed using the two-way ANOVA and Tukey's post-hoc test (ns, $p > 0.05$; *, $p < 0.05$; **, $p < 0.01$; ***, $p < 0.001$; ****, $p < 0.0001$). Interaction p values were adjusted using the Benjamini-Hochberg method to correct for false discovery rate (q). Western blotting was performed to measure ERK1/2 activation (ratio of phosphorylated ERK1/2 to total ERK1/2.
